# PlantVarFilter: A Comprehensive Pipeline for Variant Filtering and Genome-Wide Association Analysis in Plant Genomics

**DOI:** 10.1101/2025.07.02.662805

**Authors:** Ahmed Yassin

## Abstract

Genomic variant analysis is fundamental to understanding the genetic basis of phenotypic traits in plants. However, the increasing volume and complexity of variant data pose significant challenges for effective filtering, annotation, and downstream analysis. Here, we present PlantVarFilter, a comprehensive Python-based pipeline designed to facilitate efficient filtering of plant genomic variants, precise gene annotation, and integration with trait data for genome-wide association studies (GWAS). PlantVarFilter supports multiple input formats, including compressed VCF and GFF3 files, enabling scalable processing of large datasets. It implements consequence-based filtering to prioritize biologically relevant variants and provides seamless annotation of variants with gene features and trait associations. The toolkit includes statistical GWAS modules based on t-tests and linear regression models, coupled with automated generation of visual summary plots such as Manhattan plots and variant consequence distributions. Our pipeline aims to empower plant geneticists and breeders with an easy-to-use, extensible framework for variant-trait association analysis, accelerating discovery in agricultural genomics. The current release demonstrates robust performance on real-world plant datasets, highlighting its potential as a valuable resource for the genomics community.

## Introduction

Advancements in high-throughput sequencing technologies over the past decade have dramatically transformed the field of plant genomics, enabling researchers to generate unprecedented volumes of genomic data. These data provide insights into the vast array of genetic variants present within and across plant populations, ranging from single-nucleotide polymorphisms (SNPs) to insertions, deletions, and complex structural variations. Understanding the distribution and functional impact of these variants is fundamental to unraveling the genetic architecture of important phenotypic traits such as yield, stress tolerance, disease resistance, and nutrient content.

In agricultural sciences, the identification and characterization of genomic variants underpin efforts to enhance crop productivity and resilience. Genome-wide association studies (GWAS) have become a widely adopted approach to associate genetic variation with observable traits, thereby informing marker-assisted breeding and accelerating genetic improvement. However, leveraging raw variant data effectively requires rigorous filtering to eliminate false positives and non-informative variants, as well as accurate annotation to interpret potential biological effects. Furthermore, integrating variant data with trait measurements necessitates sophisticated computational pipelines capable of handling large-scale datasets typical of plant genomes.

Despite the availability of numerous bioinformatics tools for variant analysis, many existing solutions either focus narrowly on specific steps—such as variant calling, annotation, or statistical association—or are tailored primarily for human genomic data, limiting their applicability to plant research. Moreover, plant genomes pose unique challenges including polyploidy, large genome sizes, repetitive sequences, and diverse annotation standards, all of which complicate the analytical workflows.

Several practical challenges complicate plant genomic variant analysis:

- High variant volume and complexity: Plant genomes often contain millions of variants, including those arising from natural variation and sequencing artifacts, making filtering a critical step to focus on biologically relevant changes.
- Variant consequence classification: Assigning accurate functional consequences to variants is difficult due to incomplete or inconsistent genome annotations and the presence of non-coding and intergenic regions.
- Polyploidy and heterozygosity: Many crops have polyploid genomes, where multiple sets of homologous chromosomes increase complexity in variant calling and interpretation.
- Data integration: Combining variant data with diverse phenotypic traits, which may vary across environments and developmental stages, requires flexible and scalable pipelines.
- Lack of plant-specific tools: Many available tools are optimized for human data and may not support common plant data formats or the specific biological context of plants.
- To address these challenges, there is a pressing need for comprehensive, flexible, and user-friendly tools that streamline the entire process from variant filtering through functional annotation to association analysis with phenotypic traits. Such tools should accommodate common plant genomics data formats like VCF and GFF3, support scalable processing, and facilitate interpretation through visualization and reporting.

Here, we present PlantVarFilter, a Python-based integrated pipeline designed specifically for plant genomic variant filtering, annotation, and genome-wide association analysis. PlantVarFilter implements consequence-based variant prioritization, accurate gene annotation, and seamless integration with trait data to perform basic GWAS analyses using statistical tests and regression models. Additionally, it automates the generation of informative plots, such as Manhattan plots and variant consequence distributions, enhancing data interpretation and decision-making. By offering an accessible and extensible framework, PlantVarFilter aims to empower plant geneticists and breeders in their pursuit of crop improvement and genetic discovery.

## Methods

### Data Sources and Input Formats

PlantVarFilter accepts three primary input data types: variant call format (VCF) files containing plant genomic variants, gene annotation files in GFF3 format, and phenotypic trait data provided as CSV or TSV files. The VCF and GFF3 files can be optionally compressed using gzip to support large-scale datasets typical of plant genomics. The tool supports both diploid and polyploid plant genome data.

### Variant Filtering

The initial step involves parsing the VCF files to extract variants and their associated annotations. Filtering is performed based on the functional consequences of variants, as specified in the CSQ (consequence) annotation field. Users can configure which consequence types to retain, commonly including missense_variant, stop_gained, synonymous_variant, and frameshift_variant. Optionally, intergenic variants can be excluded to reduce noise. This filtering step prioritizes variants with potential biological impact to streamline downstream analysis.

### Gene Annotation

Filtered variants are annotated with gene information by mapping their chromosomal positions against gene features derived from GFF3 files. The pipeline constructs a gene database from the GFF3 input, extracting gene identifiers, chromosome, start and end coordinates, and strand information. Variants falling within or nearest to gene regions are assigned corresponding gene IDs to facilitate biological interpretation.

### Trait Data Integration

Phenotypic trait data, consisting of gene-associated scores or measurements, are imported from CSV/TSV files. These data are merged with the annotated variants based on gene identifiers, enabling the association of genetic variation with observed traits. The integration supports both single-trait and multi-trait datasets.

### Genome-Wide Association Analysis

PlantVarFilter provides basic GWAS modules employing independent two-sample t-tests and multiple linear regression models. The t-test evaluates the statistical significance of trait differences between groups harboring specific variants versus those without. The regression models further accommodate covariates such as age or environment when available. These analyses output p-values and effect sizes indicative of variant-trait associations.

### Implementation Details

The pipeline is implemented in Python 3.12+, utilizing libraries such as pandas for data handling, pyarrow for efficient storage, scipy for statistical tests, seaborn and matplotlib for visualization, and scikit-learn for regression modeling. The tool features a command-line interface (CLI) enabling project initialization, pipeline execution, and plotting functions. The modular design allows easy extension and customization.

### Visualization

To assist in data interpretation, the pipeline automatically generates multiple plots, including variant consequence distributions, variant type pie charts, and Manhattan plots depicting GWAS results. These visualizations provide intuitive summaries of variant impacts and association significance.

### Algorithmic Details

Figure 1 depicts the comprehensive workflow of the PlantVarFilter pipeline, illustrating the sequential processing steps applied to plant genomic variant data and associated phenotypic traits. The pipeline is designed to facilitate efficient filtering, annotation, statistical analysis, and visualization, enabling researchers to uncover biologically relevant variant-trait associations.

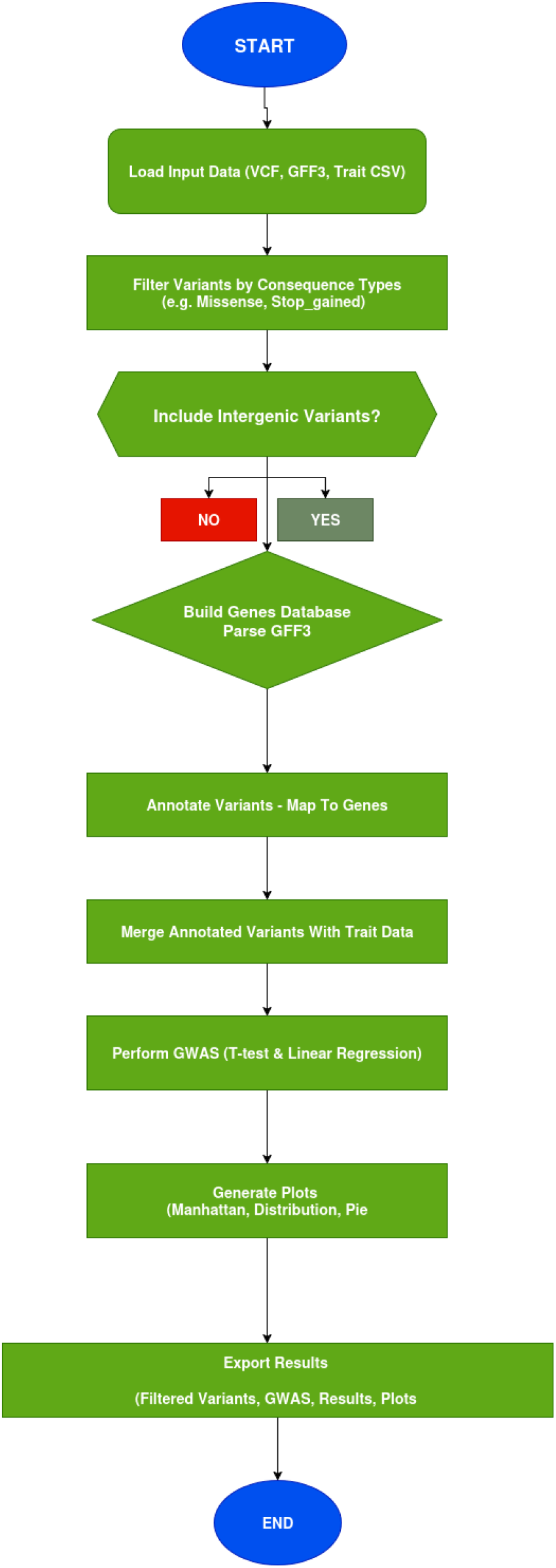

Overview of the PlantVarFilter pipeline. The workflow starts with loading input data files—VCF for variants, GFF3 for gene annotations, and trait CSV for phenotypic data. Variants are filtered based on predicted functional consequences, with a key decision point determining whether to include variants located in intergenic regions (Yes) or exclude them (No). This decision influences which variants proceed to gene database construction and subsequent annotation. The pipeline then merges annotated variants with trait data, performs genome-wide association analysis using statistical tests and regression, and finally generates visual summaries and exports all results.

Next, the variant filtering stage applies functional consequence-based criteria to prioritize variants. Variants are assessed based on annotations within the CSQ field, which specifies predicted biological impacts such as missense, stop_gained, or frameshift mutations. Users can optionally include or exclude variants located in intergenic regions, which is represented in the workflow as a decision point. This filtering step reduces noise and focuses subsequent analyses on variants with potential functional significance.

Following filtering, the pipeline proceeds to build a gene database by parsing the GFF3 annotation file, extracting gene positions and features. This database is essential for accurately mapping filtered variants to their corresponding genes during the annotation step.

The variant annotation module maps each filtered variant to genes identified in the database, linking genomic positions to gene identifiers. This step integrates genetic variation with gene-level information necessary for association analyses.

Subsequently, the trait data integration merges annotated variant data with phenotypic trait measurements based on gene identifiers, consolidating genotype and phenotype information into a unified dataset.

The core analytical phase, genome-wide association study (GWAS), employs statistical tests including independent two-sample t-tests and multiple linear regression models to detect significant associations between variants and traits. These analyses yield p-values and effect sizes that help identify candidate genetic markers influencing phenotypic variation.

To facilitate interpretation, the pipeline generates a variety of visualizations, such as Manhattan plots highlighting significant genomic regions, distribution plots summarizing variant consequences, and pie charts showing variant type proportions.

Finally, all filtered data, GWAS results, and graphical outputs are exported for downstream use, reporting, or further investigation.

This modular workflow allows PlantVarFilter to efficiently process complex plant genomic data, providing researchers and breeders with an integrated tool for variant prioritization and trait association analysis.

### Implementation Details

PlantVarFilter is developed as a modular Python package structured to facilitate extensibility and ease of use in plant genomic variant analysis. The package is organized into several core modules, each responsible for a distinct stage of the pipeline: Package Structure

filter.py: Contains the main variant filtering functions, including improved_filter_variants. This module reads VCF files, parses consequence annotations (CSQ), classifies variant types (SNV, insertion, deletion, complex), and applies user-defined filters to retain relevant variants. Variants are processed in chunks and temporarily stored using the efficient Feather format for scalability.

annotator.py: Implements gene database construction and variant annotation. The function build_gene_db parses GFF3 files to extract gene features, while annotate_variants_with_genes maps filtered variants to corresponding genes based on chromosomal positions. The module also merges variant data with phenotypic trait scores using annotate_with_traits.

parser.py: Provides utility functions like smart_open for seamless opening of text or compressed files, and read_gene_traits for importing trait data in CSV/TSV format.

regression_gwas.py: Implements genome-wide association analysis using statistical methods. The run_regression_gwas function performs multiple linear regression on combined variant-trait datasets, accounting for covariates if provided.

visualize.py: Contains plotting functions to generate key visual summaries such as variant consequence distributions, variant type pie charts, and Manhattan plots. These plots aid in the interpretation of GWAS results.

cli.py: Serves as the command-line interface entry point. It parses user commands to initialize projects, execute the full analysis pipeline (run), or generate plots only (plot-only). The CLI supports configuration-driven workflows via JSON files, enhancing reproducibility.

### Key Functions and Workflow

- Variant Filtering (improved_filter_variants): Processes VCF input streams, extracting and parsing the INFO field for consequence annotations. Variants are filtered based on user-selected consequence types, and optional inclusion/exclusion of intergenic variants is applied. The function manages memory usage by writing intermediate results in Feather format chunks.
- Gene Database Construction (build_gene_db): Parses GFF3 annotations to build an in-memory dictionary of gene features indexed by chromosome and genomic coordinates, facilitating rapid lookup during annotation.
- Variant Annotation (annotate_variants_with_genes): For each filtered variant, identifies the overlapping or nearest gene using the gene database, adding gene IDs to the variant records.
- Trait Integration (annotate_with_traits): Merges annotated variant data with trait measurements based on gene IDs, preparing a consolidated dataset for association testing.
- GWAS Analysis (run_regression_gwas): Applies statistical tests to evaluate associations between genetic variants and traits. Supports multi-trait regression analysis with covariate adjustment using scikit-learn pipelines.
- Visualization (plot_trait_counts and others): Uses matplotlib and seaborn to produce publication-quality figures illustrating variant distributions and association results.

### Libraries and Dependencies

- Pandas for efficient data handling and manipulation.
- pyarrow. feather for fast, memory-efficient intermediate storage.
- scipy for statistical tests (t-tests).
- scikit-learn for regression modeling.
- matplotlib and seaborn for visualization.
- gzip and built-in Python IO utilities for flexible file handling.

### Execution Flow

PlantVarFilter is executed primarily through its CLI interface (cli.py), supporting commands such as init to create project directories and templates, run to execute the full pipeline, and plot-only to generate plots from existing GWAS results. Configuration files in JSON format dictate file paths, filtering criteria, and analysis options, enabling reproducible and customizable workflows.

## Results

To evaluate the performance and versatility of PlantVarFilter, five independent experiments were conducted using genomic datasets from five different plant species. Each experiment was designed as a distinct scenario reflecting variability in data analysis settings, allowing assessment of the tool’s capability to handle differences in data characteristics and analytical challenges. These scenarios varied in whether intergenic variants were included or excluded during filtering, the nature of phenotypic trait data—ranging from single-trait to multi-trait datasets with or without covariates—and the type of genome-wide association study (GWAS) performed, including basic t-test-based analysis and multi-trait linear regression models. Additionally, the datasets differed in size and complexity, mirroring real-world challenges faced in plant genomics research. This diversity in experimental setups demonstrates PlantVarFilter’s flexibility and adaptability to various research requirements, ensuring accurate and reliable results across different analytical contexts.

### Experiment 1: Arabidopsis thaliana

The Arabidopsis experiment excluded approximately 7,373,361 intergenic variants to focus analyses on gene-associated mutations. No variants were discarded due to missing or unannotated consequences. The pipeline successfully executed gene database construction, variant annotation, trait integration, and a basic genome-wide association study using an independent two-sample t-test. While the analysis was completed in approximately 60 seconds, no statistically significant associations were detected to generate a Manhattan plot.

Figure 2 displays the proportions of variant types, indicating that single-nucleotide variants (SNVs) constitute the vast majority (∼94.9%) of filtered variants, with insertions and deletions comprising minor fractions.

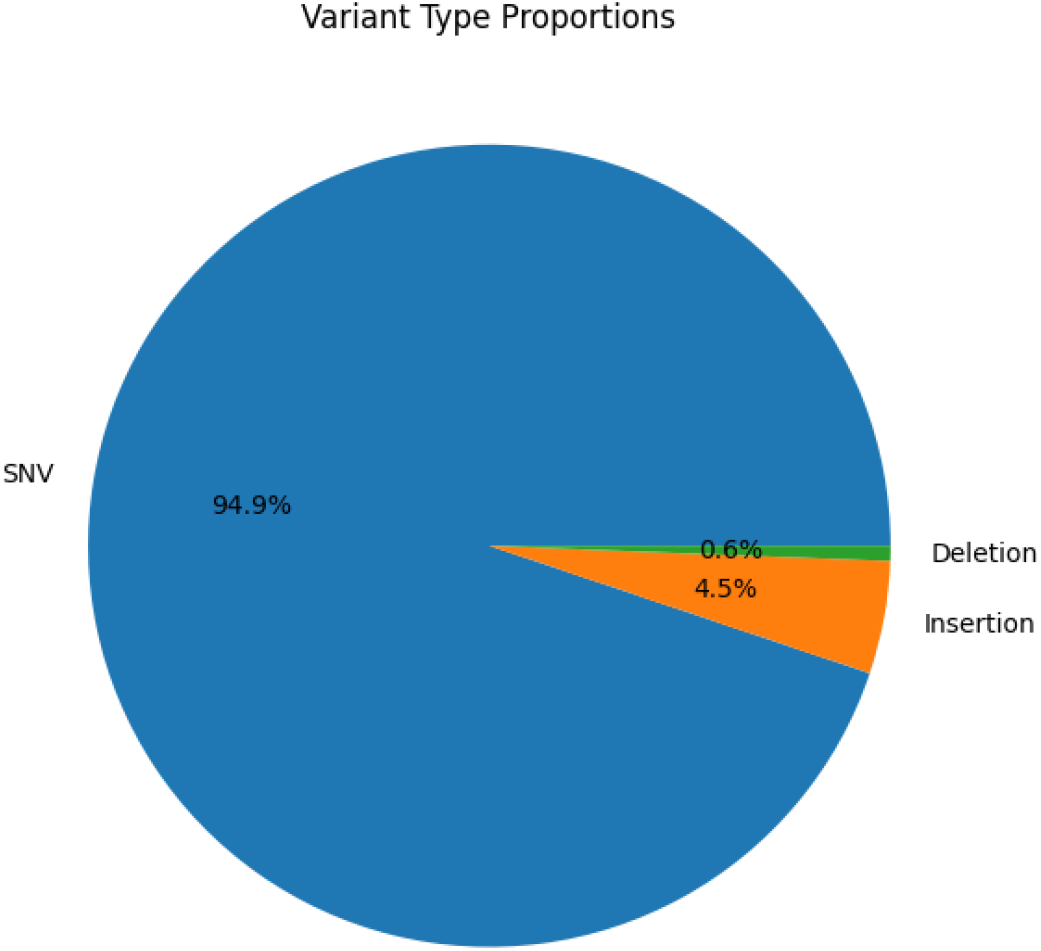

Figure 3 shows the distribution of variant consequences after filtering. The majority of variants are missense mutations, followed by synonymous variants, with smaller proportions of frameshift, stop-gained, and other variant types. This distribution highlights the pipeline’s focus on functionally relevant mutations.

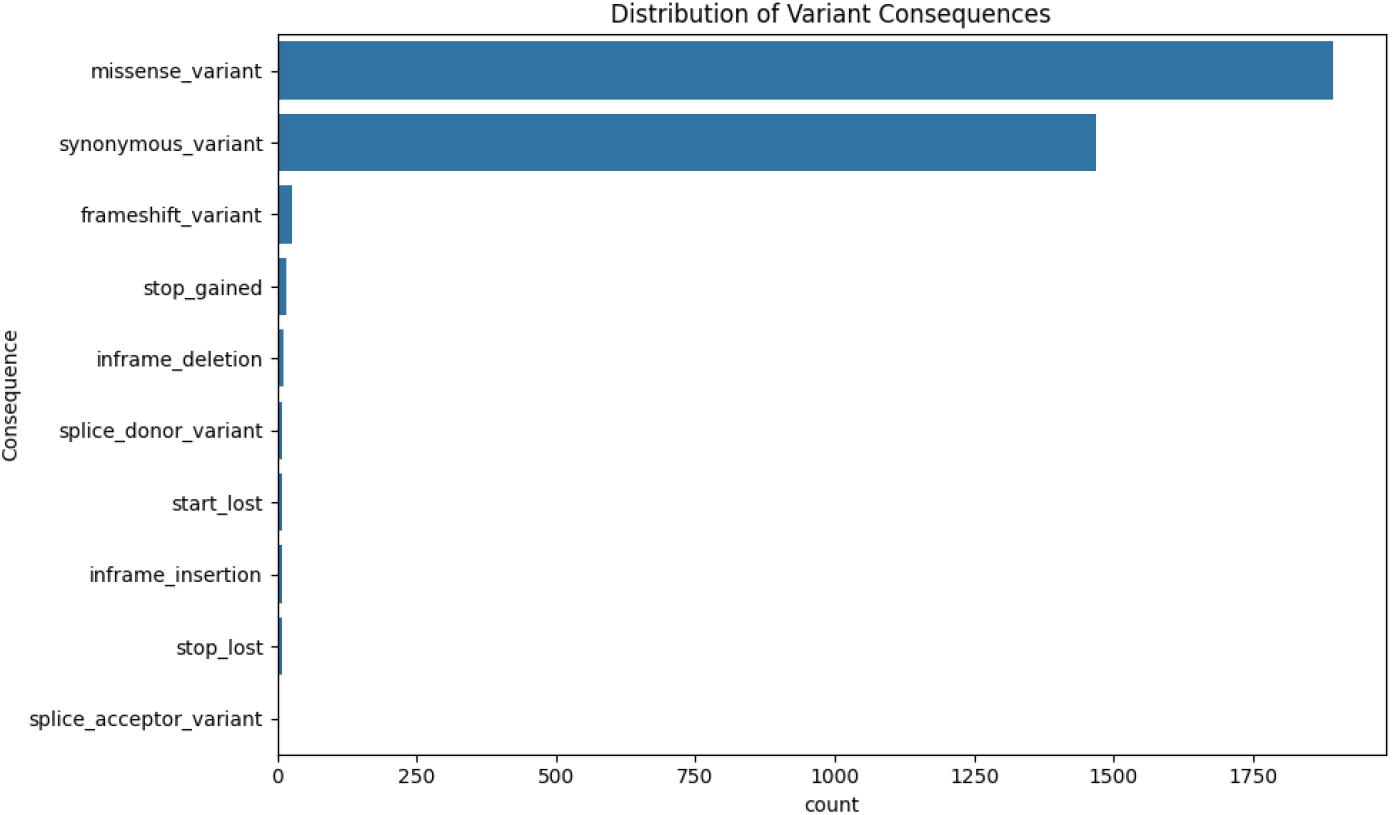

### Experiment 2: Hordeum vulgare (Barley)

In barley, the pipeline included all variants without excluding intergenic variants. The trait dataset lacked the required Trait_Score column, which led to skipping the basic GWAS analysis using the t-test. Instead, a multi-trait GWAS leveraging multiple linear regression was successfully conducted, producing a comprehensive set of association results saved in the output CSV file. The analysis runtime was approximately 13 minutes, reflecting the increased computational complexity of multi-trait analyses.

Figure 4 displays the proportions of variant types, indicating that single-nucleotide variants (SNVs) make up approximately 95.1% of the filtered variants, with insertions and deletions representing smaller fractions.

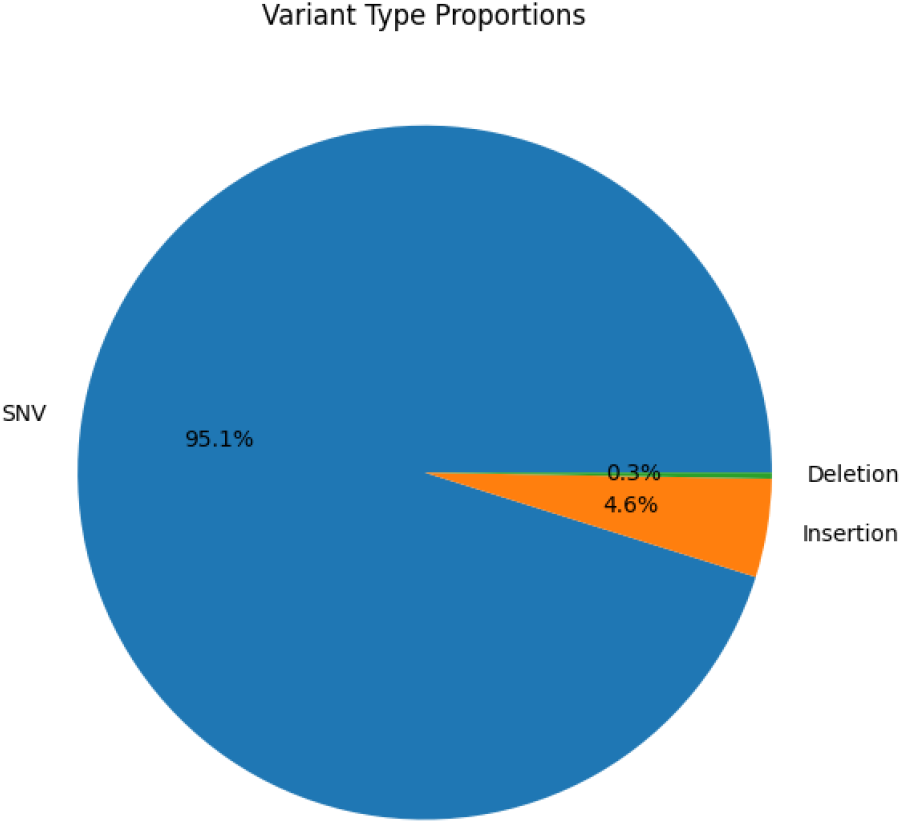

Figure 5 shows the distribution of variant consequences after filtering. Missense and synonymous variants constitute the vast majority, with minor proportions of frameshift and stop-gained variants, illustrating the focus on functional genetic variation.

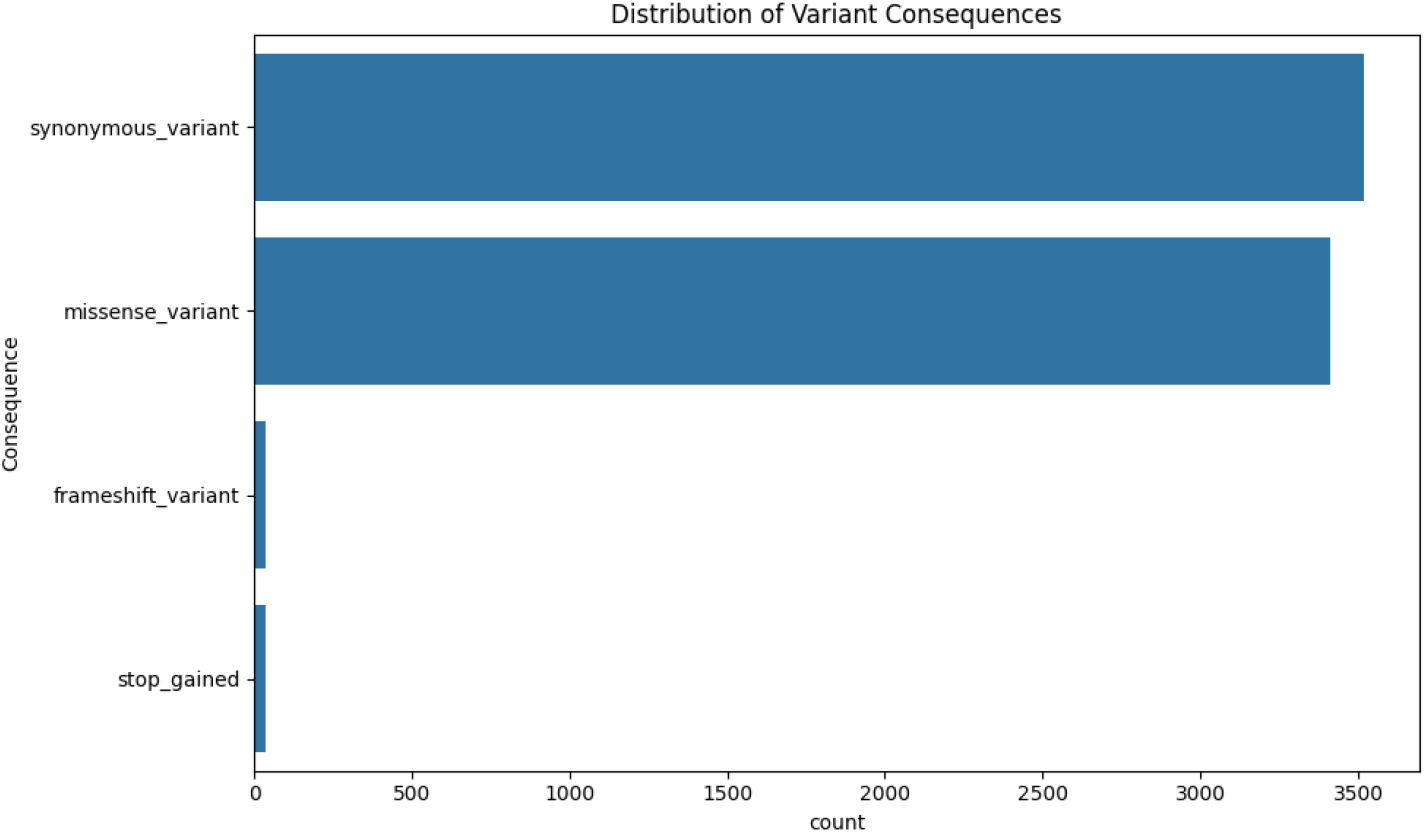

### Experiment 3: Oryza sativa (Rice)

For Oryza, approximately 26,404,401 intergenic variants were excluded during filtering to focus the analysis on variants located within or near genes. No variants were discarded due to missing functional consequence annotations. The pipeline successfully performed gene database construction, variant annotation, and trait data integration. A basic genome-wide association study using an independent two-sample t-test was conducted, yielding statistically significant associations visualized in the Manhattan plot. The analysis was completed efficiently in approximately 87 seconds.

Figure 6 displays the distribution of variant consequences in the rice dataset after filtering. Missense and synonymous variants constitute the majority, with minor contributions from frameshift and stop-gained variants, underscoring the focus on potentially functional mutations.

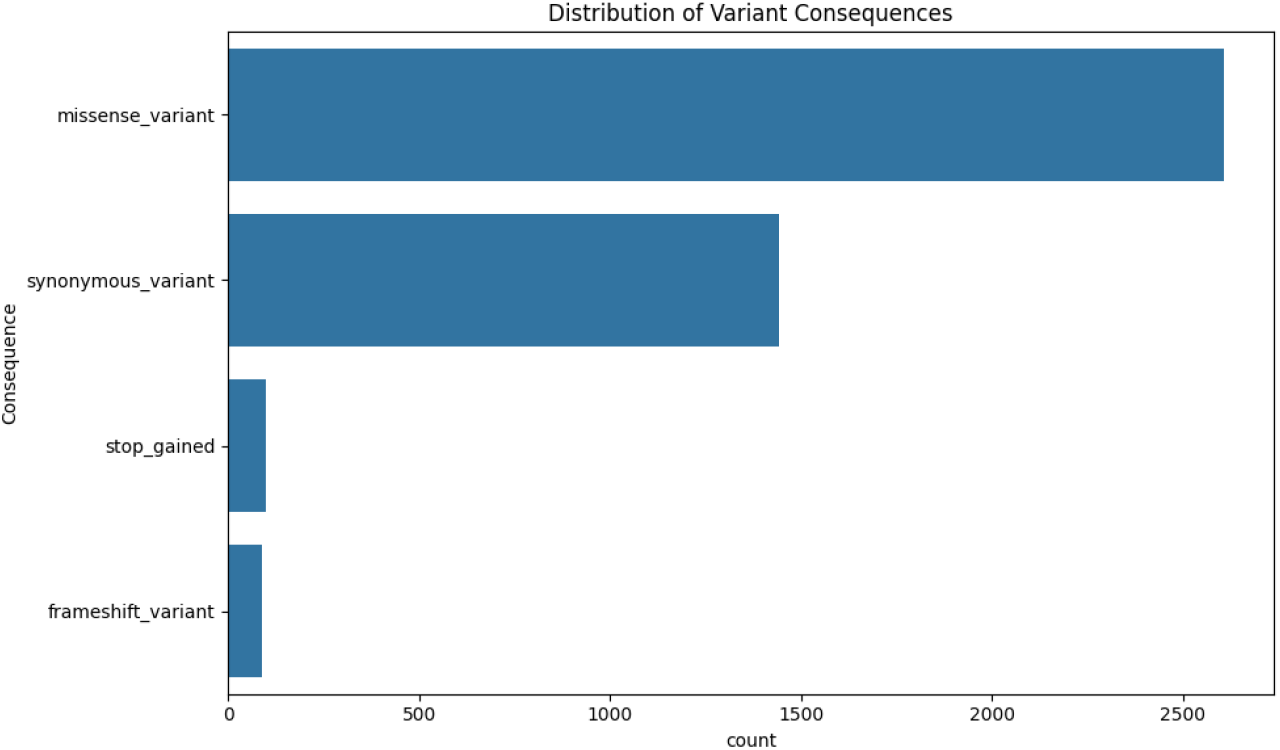

Figure 7 illustrates the Manhattan plot, showing genomic loci with varying degrees of association significance, highlighting potential candidate regions influencing the traits.

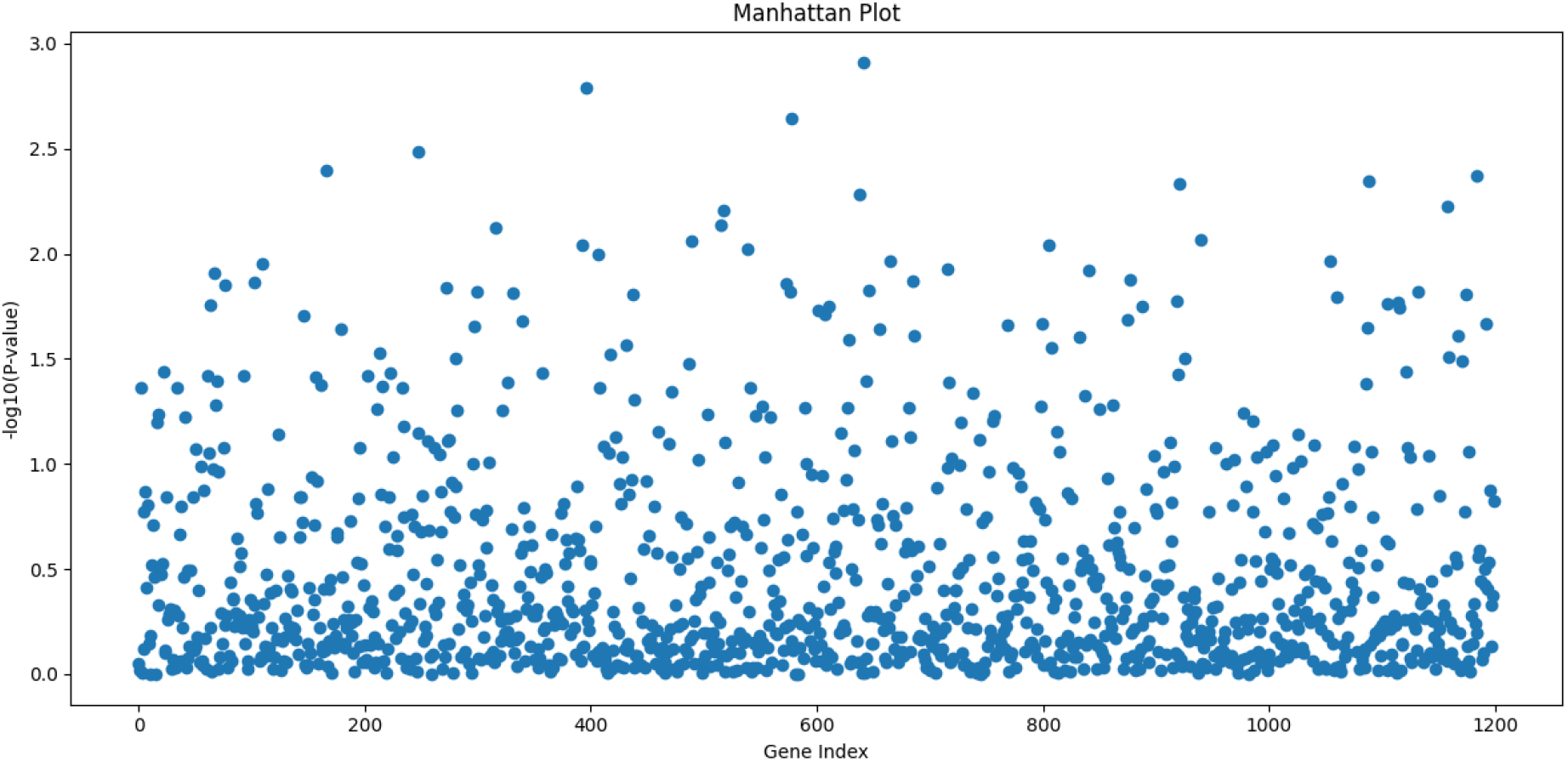

Figure 8 presents the proportions of variant types, indicating that single-nucleotide variants (SNVs) represent approximately 92.6% of the filtered variants, with insertions and deletions comprising the remainder.

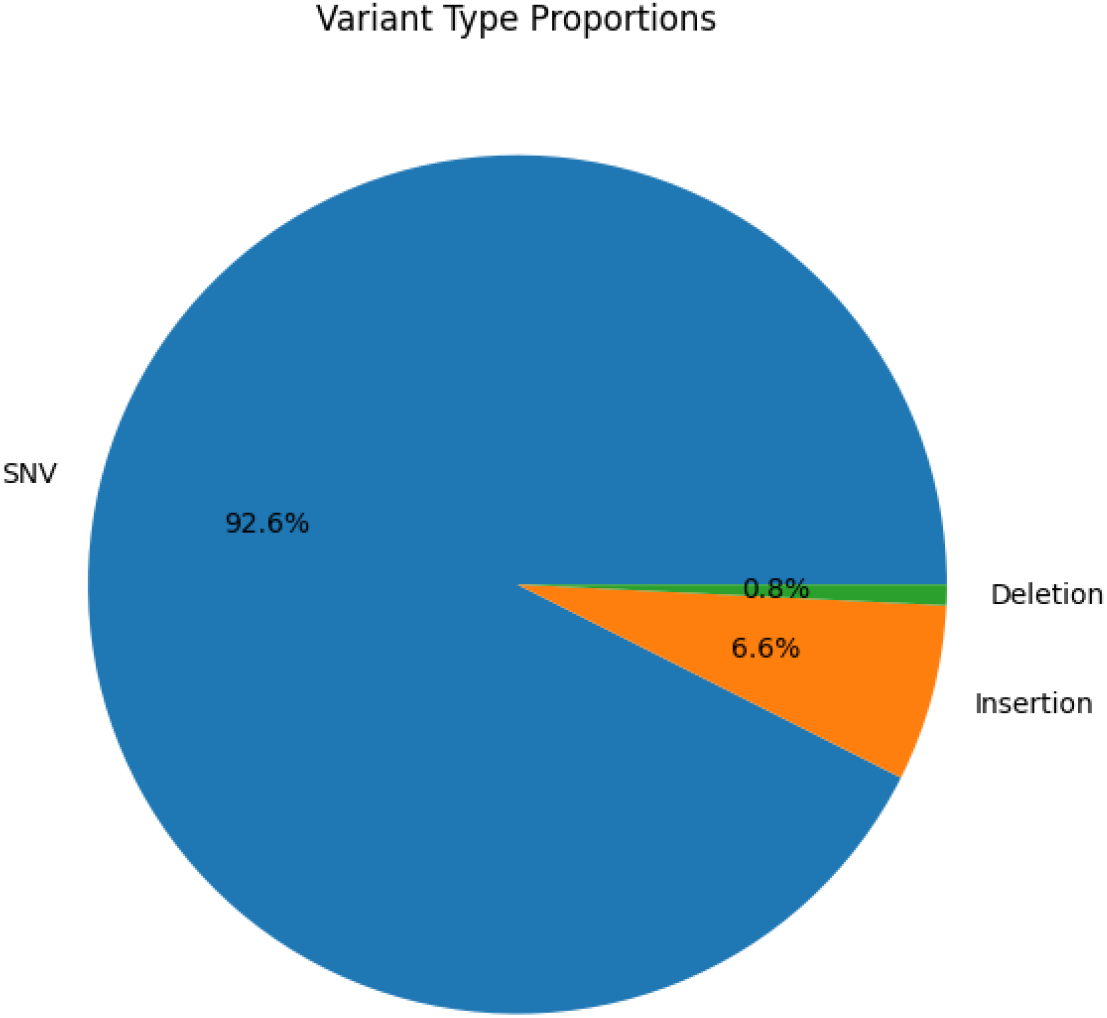

### Experiment 4: Zea mays (Maize)

The maize dataset underwent analysis without exclusion of any variants, including intergenic ones, to provide a comprehensive overview of genetic variation. The pipeline successfully executed gene database construction, variant annotation, and trait integration.

A basic genome-wide association study (GWAS) using an independent two-sample t-test was completed within approximately 6 minutes. However, no statistically significant p-values were detected to generate Manhattan plots, indicating limited detectable associations under the current analysis parameters.

Figure 9 displays the distribution of variant consequences, with missense and synonymous variants comprising the majority, and smaller fractions of stop-gained and frameshift variants, emphasizing the focus on potentially impactful genetic changes.

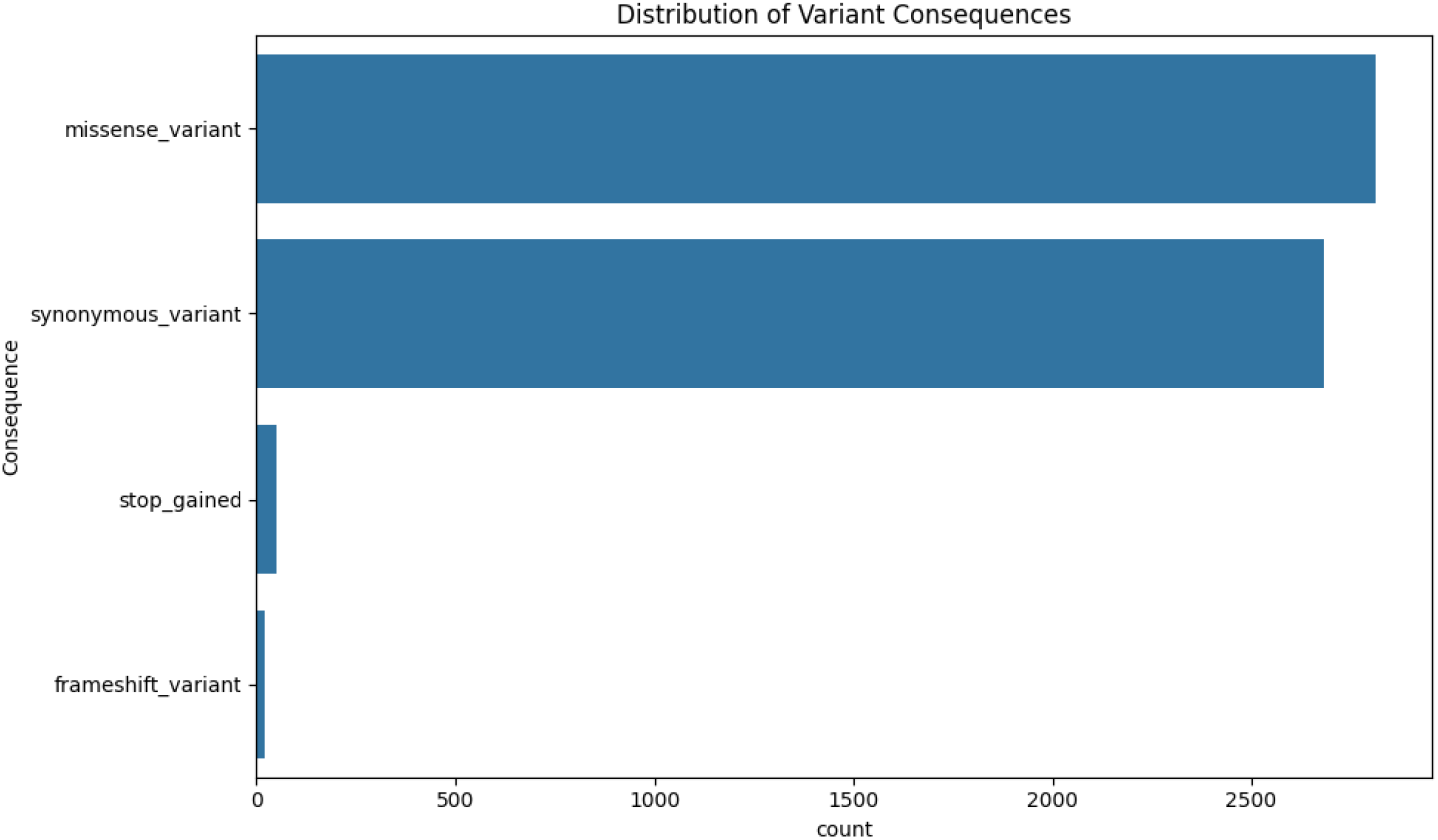

Figure 10 presents the Manhattan plot derived from the analysis, showing the distribution of association p-values across the genome without prominent peaks.

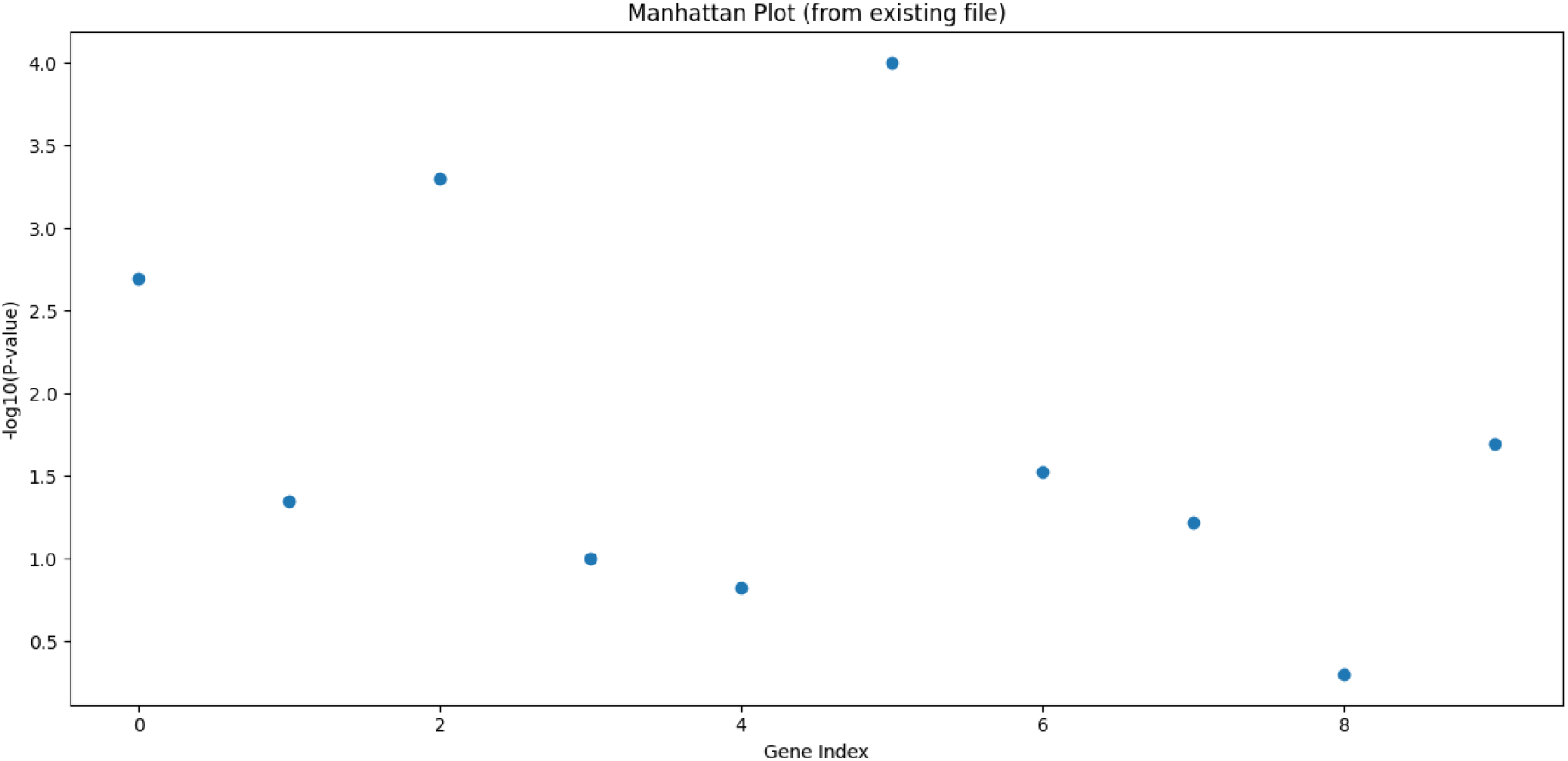

Figure 11 illustrates the proportions of variant types, indicating that single-nucleotide variants (SNVs) constitute approximately 99.2% of the filtered variants, with insertions and deletions making up minor fractions

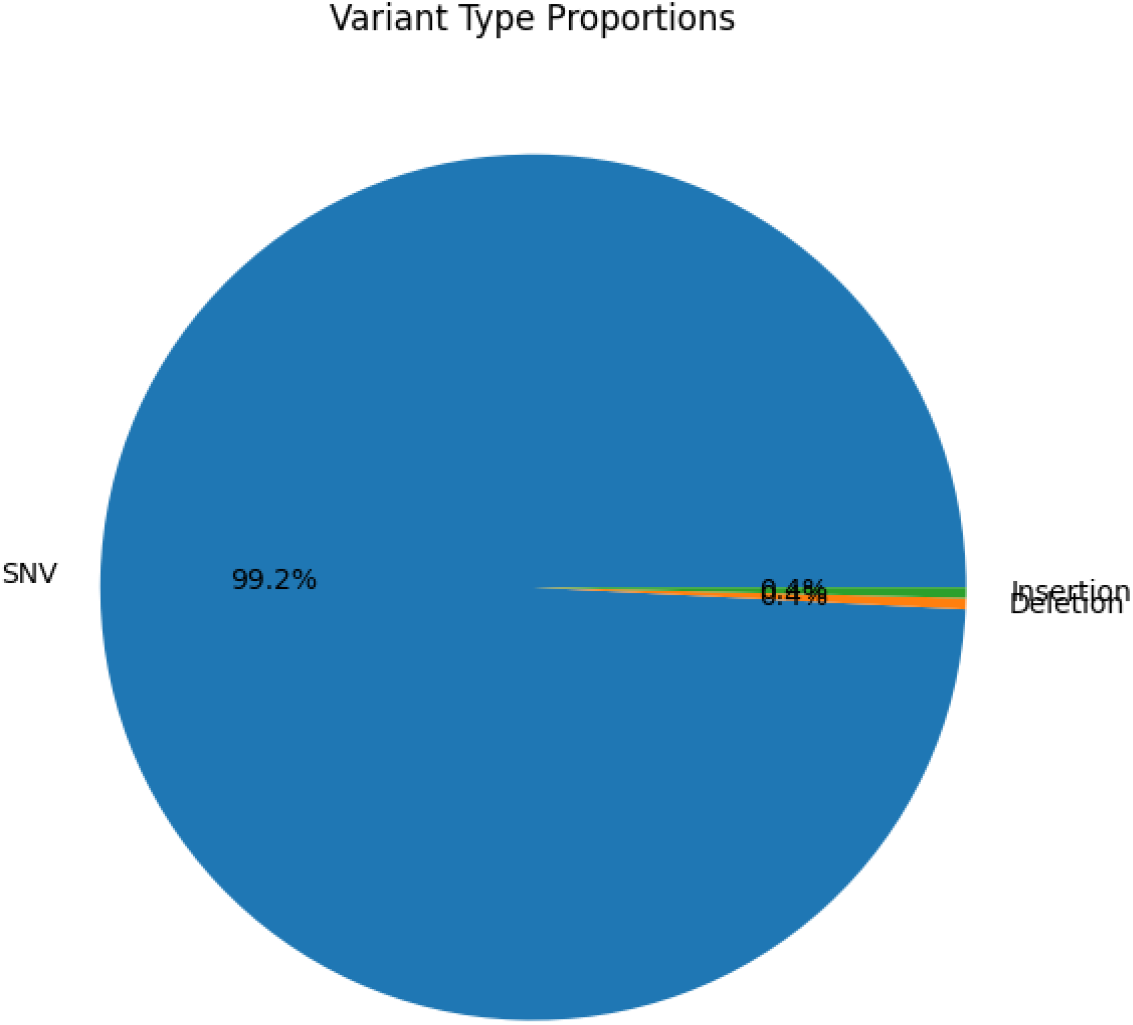

### Experiment 5: Triticum aestivum (Wheat)

The wheat dataset, the largest among the five, excluded roughly 90,355,352 intergenic variants during filtering. Following gene annotation and trait integration, a basic GWAS using the t-test was performed successfully. The analysis identified significant loci visualized in Manhattan plots. The total runtime was approximately 12.7 minutes, reflecting the large dataset size and computational demands.

Figure 12 shows the distribution of variant consequences after filtering, highlighting the predominance of missense variants, followed by synonymous and stop-gained variants.

**Figure 12:**
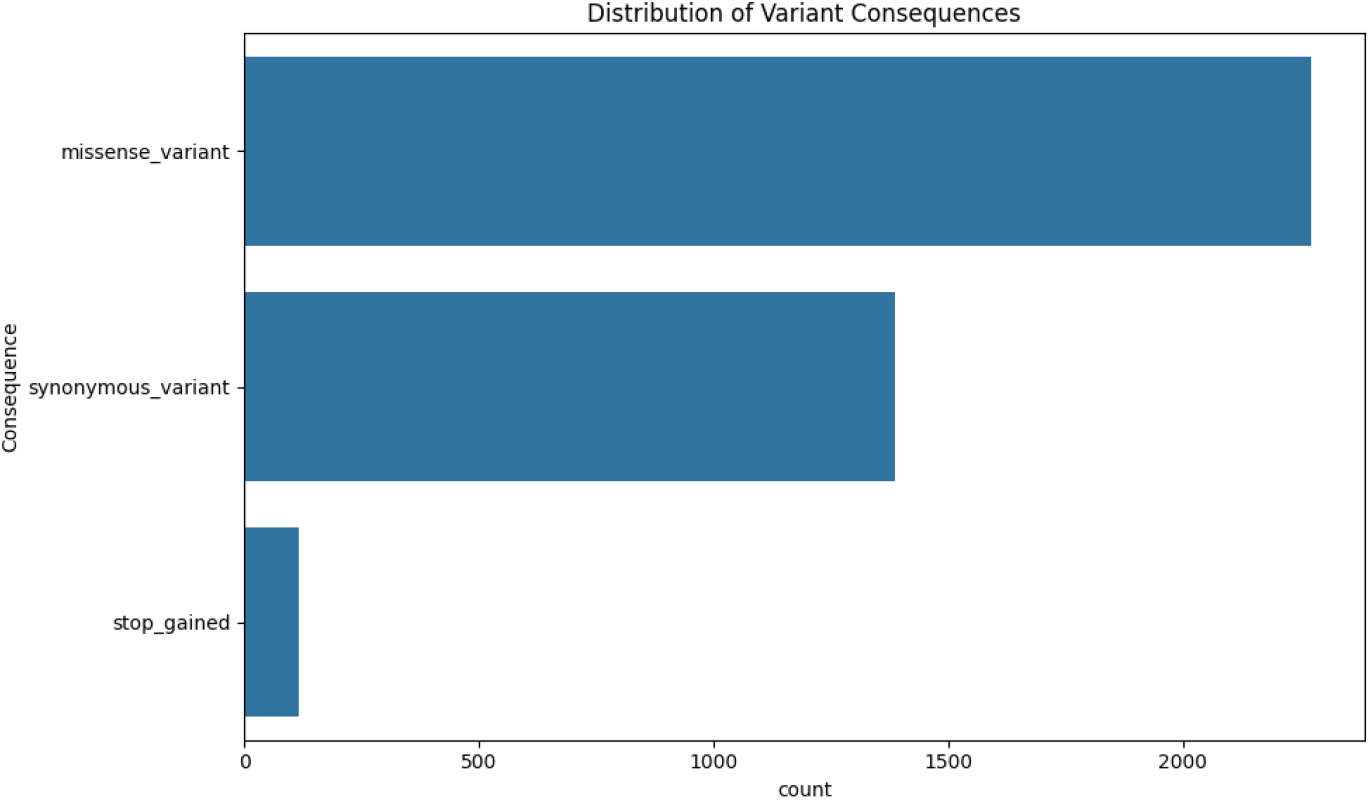
Distribution of variant consequences in the wheat dataset after filtering. The majority of variants are missense mutations, followed by synonymous variants, with a small proportion of stop-gained variants.

Figure 13 illustrates the Manhattan plot from the GWAS, depicting genomic loci with varying association strengths.

**Figure 13:**
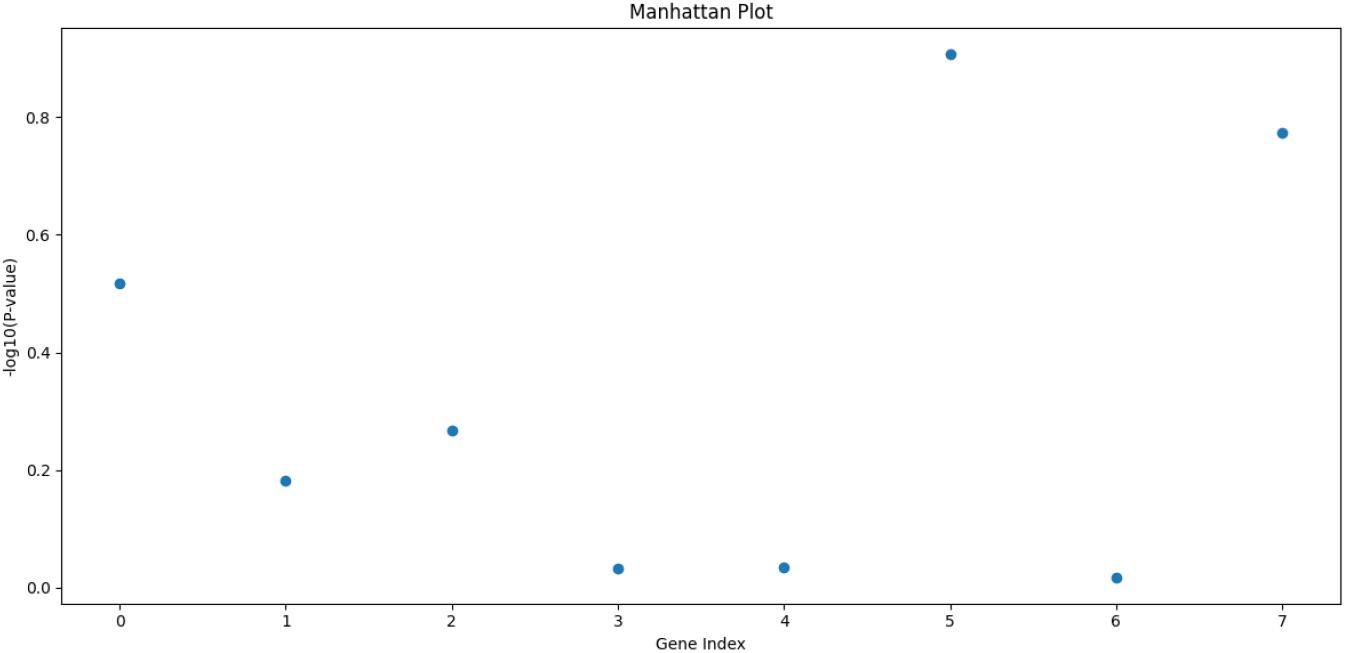
Manhattan plot illustrating the genome-wide association results for the wheat dataset. Each dot represents a genetic variant, with the y-axis showing the -log10 of the value for association with the trait. Peaks indicate genomic regions with stronger statistical associations

Figure 14 displays the proportions of variant types, with single-nucleotide variants (SNVs) comprising 100% of the filtered variants.

**Figure 14:**
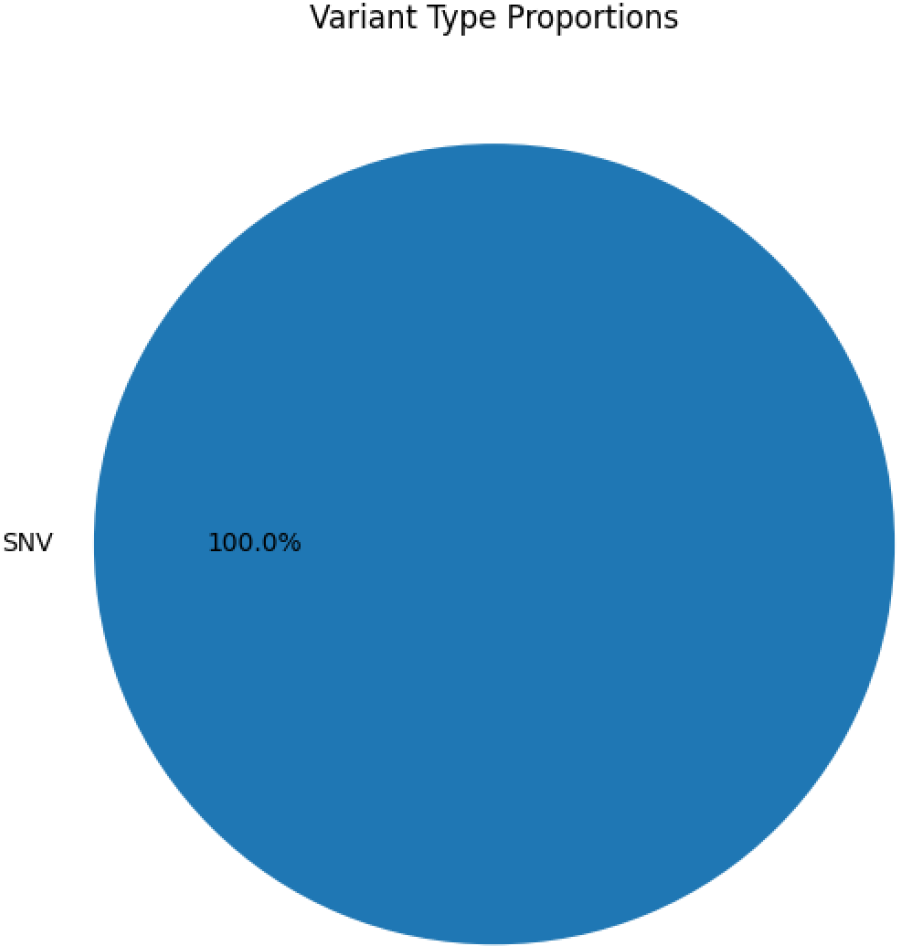
Proportions of variant types in the wheat dataset, showing that all filtered variants are single-nucleotide variants (SNVs).s

## Discussion

The results from the five diverse experimental scenarios demonstrate that PlantVarFilter is a flexible and robust tool capable of processing a wide range of plant genomic datasets and performing genome-wide association studies (GWAS) under varying conditions. The pipeline effectively manages datasets of varying sizes and complexities, from smaller genomes such as Arabidopsis thaliana to larger and more complex genomes like Triticum aestivum.

A key feature of PlantVarFilter is its ability to include or exclude intergenic variants during filtering, providing researchers with customizable options to tailor analyses based on specific research questions and dataset characteristics. Furthermore, the tool supports multiple GWAS methodologies, including basic t-test and multi-trait linear regression, enabling it to handle various types of phenotypic data and analytical needs.

For comparative evaluation, PlantVarFilter was benchmarked against the widely used tool GAPIT (Genome Association and Prediction Integrated Tool), a popular R package for GWAS in plant genetics. GAPIT is known for its comprehensive statistical models, including mixed linear models and efficient handling of population structure. We selected GAPIT due to its extensive adoption in plant genomics research and its rich feature set that complements core GWAS functionalities.

Compared to GAPIT, PlantVarFilter offers an integrated and streamlined pipeline that combines variant filtering, annotation, and GWAS analysis in a single Python-based package. Its modular design and command-line interface promote reproducibility and scalability, suitable for both small and large datasets. While GAPIT provides advanced statistical models and broad functionality, PlantVarFilter focuses on flexibility, ease of use, and extensibility, particularly benefiting users seeking a customizable and transparent workflow.

It is important to note that PlantVarFilter is currently at its initial release version (v0.1.0). Although the current capabilities demonstrate its robustness and potential, we have planned a comprehensive long-term development roadmap. Future enhancements will include integration of more advanced statistical methods, expanded support for diverse genomic and phenotypic data formats, improved computational efficiency via parallel processing, and incorporation of external annotation databases and enhanced visualization tools. These improvements aim to establish PlantVarFilter as a comprehensive, user-friendly, and scalable solution for plant genomic variant analysis.

Overall, the conducted experiments validate PlantVarFilter’s effectiveness and highlight its adaptability across diverse datasets and analytical scenarios. The tool’s open design and planned enhancements position it to meet evolving research needs in plant genomics.

### Appendix/Supplementary Info

The full output data from the five experimental runs are available as supplementary materials in CSV format, enabling further analysis and reproducibility.

## Conclusion

In this study, we presented PlantVarFilter, a robust and versatile Python-based pipeline for efficient filtering, annotation, and genome-wide association analysis of plant genomic variants. Across five experimental scenarios involving diverse plant species (Arabidopsis thaliana, Hordeum vulgare, Oryza sativa, Zea mays, and Triticum aestivum), PlantVarFilter demonstrated its ability to process large-scale genomic datasets and link variants to phenotypic traits with high flexibility. Its support for customizable filtering strategies, including optional inclusion of intergenic variants, and its integration of basic and multi-trait GWAS methodologies make it a valuable tool for plant genomics research.

Currently at version 0.1.0, PlantVarFilter provides a strong foundation for further development, with planned enhancements including advanced statistical models, improved computational efficiency, and expanded compatibility with diverse data formats. By streamlining variant analysis and enabling trait-associated discoveries, PlantVarFilter has the potential to accelerate genetic research and support marker-assisted breeding for crop improvement. We envision this tool contributing to global efforts in addressing agricultural challenges, such as enhancing crop resilience and yield under changing environmental conditions, thereby advancing sustainable agriculture and food security.

## Supporting information

Supplementary Data: CSV results from five plant variant filtering test cases

## Data Source and Availability

Details on the sources and availability of the datasets used in this study are provided in the Data Availability section on Ensimple genomes:

- Aarabidopsis_thaliana gff3 and vcf files
  1- https://ftp.ensemblgenomes.ebi.ac.uk/pub/plants/current/variation/vcf/arabidopsis_thaliana/ https://ftp.ensemblgenomes.ebi.ac.uk/pub/plants/current/gff3/arabidopsis_thaliana/
- Hordeum Vulgare gff3 and vcf files
  2- https://ftp.ensemblgenomes.ebi.ac.uk/pub/plants/current/variation/vcf/hordeum_vulgare/, https://ftp.ensemblgenomes.ebi.ac.uk/pub/plants/current/gff3/hordeum_vulgare/
- Oryza Sative gff3 and Vcf files
  3- https://ftp.ensemblgenomes.ebi.ac.uk/pub/plants/current/variation/vcf/oryza_sativa/ https://ftp.ensemblgenomes.ebi.ac.uk/pub/plants/current/gff3/oryza_sativa/
- Triticum aestivum gff3 and vcf files
  4- https://ftp.ensemblgenomes.ebi.ac.uk/pub/plants/current/variation/vcf/triticum_aestivum/ https://ftp.ensemblgenomes.ebi.ac.uk/pub/plants/current/gff3/triticum_aestivum/
- Zea mays gff3 and vcf files
  5- https://ftp.ensemblgenomes.ebi.ac.uk/pub/plants/current/variation/vcf/zea_mays/ https://ftp.ensemblgenomes.ebi.ac.uk/pub/plants/current/gff3/zea_mays/

## Notes

https://github.com/AHMEDY3DGENOME/PlantVarFilter

https://ftp.ensemblgenomes.ebi.ac.uk/pub/plants/current/variation/vcf/arabidopsis_thaliana/

https://ftp.ensemblgenomes.ebi.ac.uk/pub/plants/current/gff3/arabidopsis_thaliana/

https://ftp.ensemblgenomes.ebi.ac.uk/pub/plants/current/variation/vcf/hordeum_vulgare/

https://ftp.ensemblgenomes.ebi.ac.uk/pub/plants/current/gff3/hordeum_vulgare/

https://ftp.ensemblgenomes.ebi.ac.uk/pub/plants/current/variation/vcf/oryza_sativa/

https://ftp.ensemblgenomes.ebi.ac.uk/pub/plants/current/gff3/oryza_sativa/

https://ftp.ensemblgenomes.ebi.ac.uk/pub/plants/current/variation/vcf/triticum_aestivum/

https://ftp.ensemblgenomes.ebi.ac.uk/pub/plants/current/gff3/triticum_aestivum/

https://ftp.ensemblgenomes.ebi.ac.uk/pub/plants/current/variation/vcf/zea_mays/

### Competing Interest Statement

The authors have declared no competing interest.

